# Modulation of Microtubule Dynamics by Monovalent Ions

**DOI:** 10.1101/2024.06.26.600768

**Authors:** Simon Fernandes, Charlotte Aumeier

## Abstract

The microtubule cytoskeleton is a dynamic network essential for many cellular processes, influenced by physicochemical factor such as temperature, pH, dimer concentration and ionic environment. In this study, we used in vitro reconstitution assays to examine the effects of four monovalent ions (Na^+^, K^+^, Cl^-^, and Ac^-^) on microtubule dynamics, uncovering distinct effects for each ion. Na^+^ was found to increase microtubule dynamicity by raising catastrophe frequency, polymerization and depolymerization speeds, ultimately reducing microtubule lifetime by 80 %. Conversely, Ac boosts microtubule nucleation and stabilizes microtubules by increasing rescue frequency and preventing breakages, resulting in longer microtubules with extended lifetimes. Cl^-^ appeared to potentiate the effects of Na^+^, while K^+^ had minimal impact on microtubule dynamic parameters. These findings demonstrate that Na^+^ and Ac^-^ have opposing effects on microtubule dynamics, with Na^+^ destabilizing and Ac^-^ stabilizing the microtubule structure. This ionic impact is mainly through modulation of tubulin-tubulin interactions rather than affecting the hydrolysis rate. In conclusion, ion identity plays a crucial role in modulating microtubule dynamics. Understanding the ionic environment is essential for microtubule-related research, as it significantly influences microtubule behavior, stability, and interactions with other proteins.

**SIGNIFICANCE STATEMENT:** The microtubule cytoskeleton is vital for cellular processes and influenced by temperature, pH, dimer concentration, and ionic environment. Understanding how these physicochemical factors regulate microtubule polymerization is crucial for elucidating microtubule dynamics and stability. Our in vitro reconstitution assays reveal that Na^+^ and Ac^-^ ions have opposing effects on microtubule dynamics. Na^+^ increases dynamicity by raising catastrophe frequency and reducing lifetime by 80 %, while Ac ^-^ enhances nucleation and stability, resulting in longer microtubules. Cl^-^ potentiates the effects of Na^+^, and K^+^ has minimal impact. Our findings highlight that ion identity crucially modulates microtubule dynamics, significantly influencing stability and interactions.

## INTRODUCTION

The microtubule cytoskeleton is a highly dynamic network, conserved throughout all eukaryotes. Composed of αβ-tubulin heterodimers, microtubules are polarized hollow tubes that form by longitudinal and lateral interactions between the dimers (Desai and Mitchison, 1997; Schulze and Kirschner, 1986). As biopolymers, microtubule formation follows the classical physicochemical rules. Temperature, pH, and dimer concentration are central factors to determine the fate of the biopolymer (Behnke, 1964; Croom et al., 1986; Desai and Mitchison, 1997). Moreover, pioneer studies using reconstituted *in vitro* assays demonstrated that the ionic composition is a major player in microtubule biochemistry impacting microtubule dynamics.

Microtubules polymerize by the addition of GTP-tubulin dimers at their ends (Valiron et al., 2001). After incorporation, GTP is hydrolyzed to GDP resulting in an unstable GDP-microtubule in comparison to the growing GTP-tubulin end that is stabilizing the structure (Hyman et al., 1992; Nogales et al., 1998). Over time the structure of the growing end becomes more irregular, tapered, reducing the local concentration of GTP-tubulin. When GTP hydrolysis catches up with the addition of new GTP-tubulin dimers, colliding with a reduction of the local GTP-tubulin density, the unstable microtubule undergo catastrophe followed by rapid depolymerization (Bowne-Anderson et al., 2015; Farmer and Zanic, 2023; Mitchison and Kirschner, 1984). Moreover, shrinking microtubules can switch back to re-growth, an event called (Mitchison and Kirschner, 1984). Neither the addition or removal of tubulin is restricted to microtubule ends; they can happen spontaneously all along the shaft (Andreu-Carbó et al., 2022b; Dimitrov et al., 2008; Schaedel et al., 2019; Triclin et al., 2021; Tropini et al., 2012). Indeed, damages along the shaft can be repaired by the incorporation of fresh GTP-tubulin dimers which can act as rescue sites (Aumeier et al., 2016). Altogether, these dynamic parameters, polymerization and depolymerization rates, along with rescue and catastrophe frequencies, govern the microtubule lifetime.

Ions finetune protein to protein interaction by changing electrostatic affinities (Sheinerman et al., 2000; Zhou and Pang, 2018). In this scenario, tubulin-tubulin interactions during polymerization are subjected to this electrostatic modulation. Once formed, microtubules exhibit electronegative charges at their surfaces that can be mitigated by the presence of ions, which, in turn, can alter interactions with microtubule associated proteins (Drechsler et al., 2019; Tuszyński et al., 2005). Moreover, ions exhibit different lyotropic properties impacting on the solubilization of proteins and, therefore, protein stability in solution (Cacace et al., 1997; Gregory et al., 2022; Kang et al., 2020; Okur et al., 2017). This ability of ions to stabilize or destabilize proteins in solution could be relevant in the context of microtubule polymerization as soluble tubulin heterodimers polymerize into an insoluble microtubule.

Early studies on microtubule polymerization addressed the role of ions in microtubule assembly. The divalent cations Magnesium (Mg^2+^) and Calcium (Ca^2+^) were rapidly identified to influence microtubule dynamics (Kuriyama and Sakai, 1974; Rosenfeld et al., 1976). Mg^2+^ binds to the tubulin GTP pocket and acts as a co-factor for GTP hydrolysis. Consequently microtubule growth is impaired when Mg^2+^ is chelated by Ethylenediamine Tetraacetic Acid (EDTA) during the polymerization reaction (Fees and Moore, 2019; Grover and Hamel, 1994; Huang et al., 1985; Martin et al., 1987; O’Brien et al., 1990). However, when increasing the concentration of Mg^2+^ beyond 10 mM destabilizes microtubules, probably by modulating the electrostatic interactions between tubulins. Ca^2+^ destabilizes microtubule integrity even more efficiently by increasing the catastrophe frequency and the depolymerization rate (O’Brien et al., 1997, 1990; Rosenfeld et al., 1976; Weisenberg and Deery, 1981). Concentrations under 1 mM Ca^2+^ already impair microtubule growth and the addition of Ethylene Glycol Tetraacetic Acid (EGTA) rescues microtubule formation by chelating Ca^2+^ (Kuriyama and Sakai, 1974). Those findings set the base for the *in vitro* microtubule research.

Monovalent ions also impact microtubule dynamics, although in comparison with divalent cations higher concentrations are needed to achieve effects. Concentrations of monovalent salts above 250 mM prevented microtubule formation and induced tubulin aggregation (Croom et al., 1986), while at physiological concentration (50-150 mM) ions such as alkali monovalent cations favoured microtubule formation in the presence of Taxol, a microtubule stabilizing drug (Suzaki et al., 1978; Wolff et al., 1996). In particular, sodium (Na^+^) or potassium (K^+^) showed an enhancement of the net microtubule formation (Wolff et al., 1996). Similarly, monovalent anions, including acetate (Ac^-^), glutamate and fluoride enhanced the net microtubule mass (Suzaki et al., 1978). However, all these experiments were performed in bulk experiments and a comprehensive investigation into the role of monovalent ions on direct dynamical features of microtubules is missing.

Here, we show how four different monovalent ions—Na^+^, K^+^, Cl^-^, and Ac^-^ —affect microtubule dynamics and stability. Na^+^ increases microtubule dynamicity by raising catastrophe frequency and enhancing polymerization and depolymerization speeds at the microtubule ends. In contrast, Ac^-^ promotes microtubule nucleation, extends microtubule length and lifetime by enhancing rescue frequency and reducing depolymerization frequency. Additionally, the effects of these salts on microtubule stability are independent of GTP hydrolysis, as they similarly influence microtubules in both GDP-tubulin and GTP-tubulin states. Our study shed light on the complex interplay of ions in microtubule assembly, providing novel insights into the mechanistic aspects of tubulin polymerization and microtubule biochemistry.

## RESULTS

### Effect of ions on microtubule bulk dynamics

We selected KCl, NaCl, KAc and NaAc as salts to distinguish how individual monovalent ions, Na^+^, K^+^, Cl^-^ and Ac^-^, as well as associative cation-anion effects impact on microtubule dynamics. Microtubules were polymerized from 15 µM tubulin using the BRB buffer in the absence or presence of the different ions at concentration of 50 mM or 100 mM. Initially, we studied the influence of these salts on net microtubule mass formation using a turbidity assay (**Fig. 1a**). In the presence of 100 mM KAc, we saw the fastest microtubule nucleation (smallest lag time, 352 sec), the fastest bulk polymerization (steepest slope, 2.07*10^−4^ a.u./sec) reaching the highest polymer mass (plateau at 0.1171 a.u.) (**Fig. 1b**). Microtubules grown in the presence of 100 mM NaAc had comparable bulk dynamics. The addition of KCl showed a turbidity profile comparable to the control condition (**Fig. 1a and 1b**). The presence of NaCl marginally decreased net microtubule formation, resulting in the lowest plateau values compared to the control. Therefore, KAc and NaAc are potent enhancers of microtubule formation, whereas NaCl might have a destabilizing effect.

**Figure 1:**
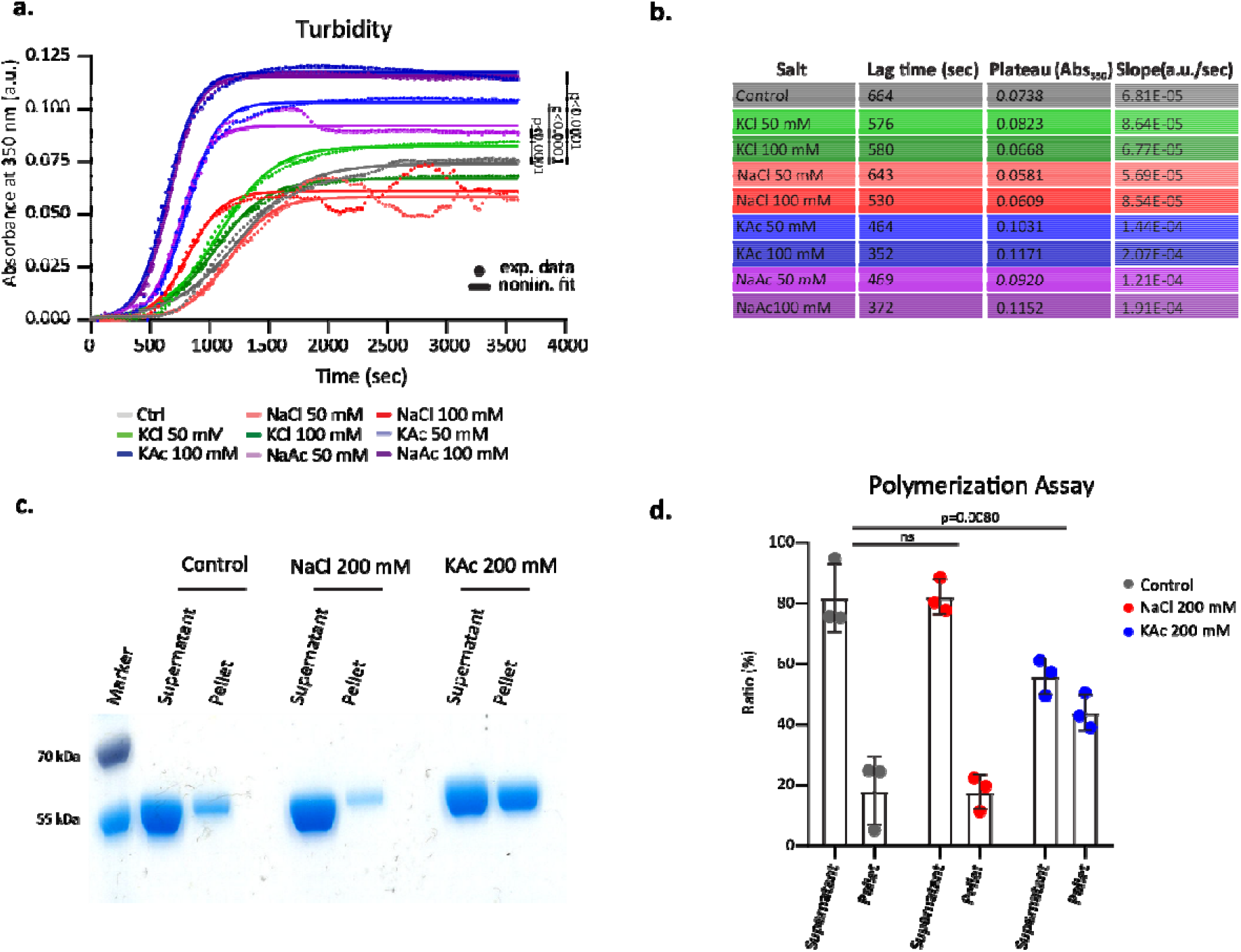
Ion effect on bulk polymerization. **a**. Representative turbidity assay of 15 μM tubulin in the presence of the indicated salts. Absorbance at 350 nm was recorded every 30 seconds at 37°C. Experimental data are represented by colored dots. A nonlinear fit (logistic growth) for each condition is plotted as a colored solid line. Statistics: one-way ANOVA. **b**. Lag time, plateau and slope were calculated from the nonlinear fit of the turbidity assay (**Fig. 1a**, see methods). **c**. Pelleting assay of 10 μM tubulin. Microtubules were polymerized for 30 minutes at 37°C in BRB80 with 1 mM GTP, 20 % (v/v) glycerol and 200 mM of the indicated salt. Representative SDS-PAGE with supernatant and pellets after 30 minutes polymerization at 37°C. **d**. Quantification of band intensity of **Fig. 1c**. Mean with SD, n = 3 independent experiments. Statistics: two-way ANOVA.

To validate those observation with a complementary bulk assay, we performed a pelleting assay wherein microtubules were polymerized from 10 µM tubulin for 30 min in the presence or absence of 200 mM salt. Doubling the salt concentration aimed to intensify the measured salt effects in **Fig. 1a**. In the presence of 200 mM KAc the polymerized fraction in the pellet increased by 2.5-fold compared to the control, confirming its stabilizing effect on net microtubule formation (**Fig. 1c and 1 d**). Microtubule formation in the presence of 200 mM NaCl was comparable to control (**Fig. 1d**).

### Effect of ions on single microtubule dynamics

While a bulk assay offers valuable insights, it lacks the resolution to discern whether higher polymer mass results from enhanced nucleation, polymerization, or stabilizing effects. To delve deeper into understanding the impact of the different salts on the parameters of microtubule dynamics – such as growth speed, depolymerization speed, catastrophe, and rescue frequency – we studied the dynamic parameters of single microtubules polymerized from seeds using a TIRF microscopy-based assay (**Fig. 2**). We used 0 mM to 100 mM salt concentrations, as at higher concentrations analysis of microtubule dynamics was not possible for NaCl and KAc (**Fig. 2a**).

**Figure 2:**
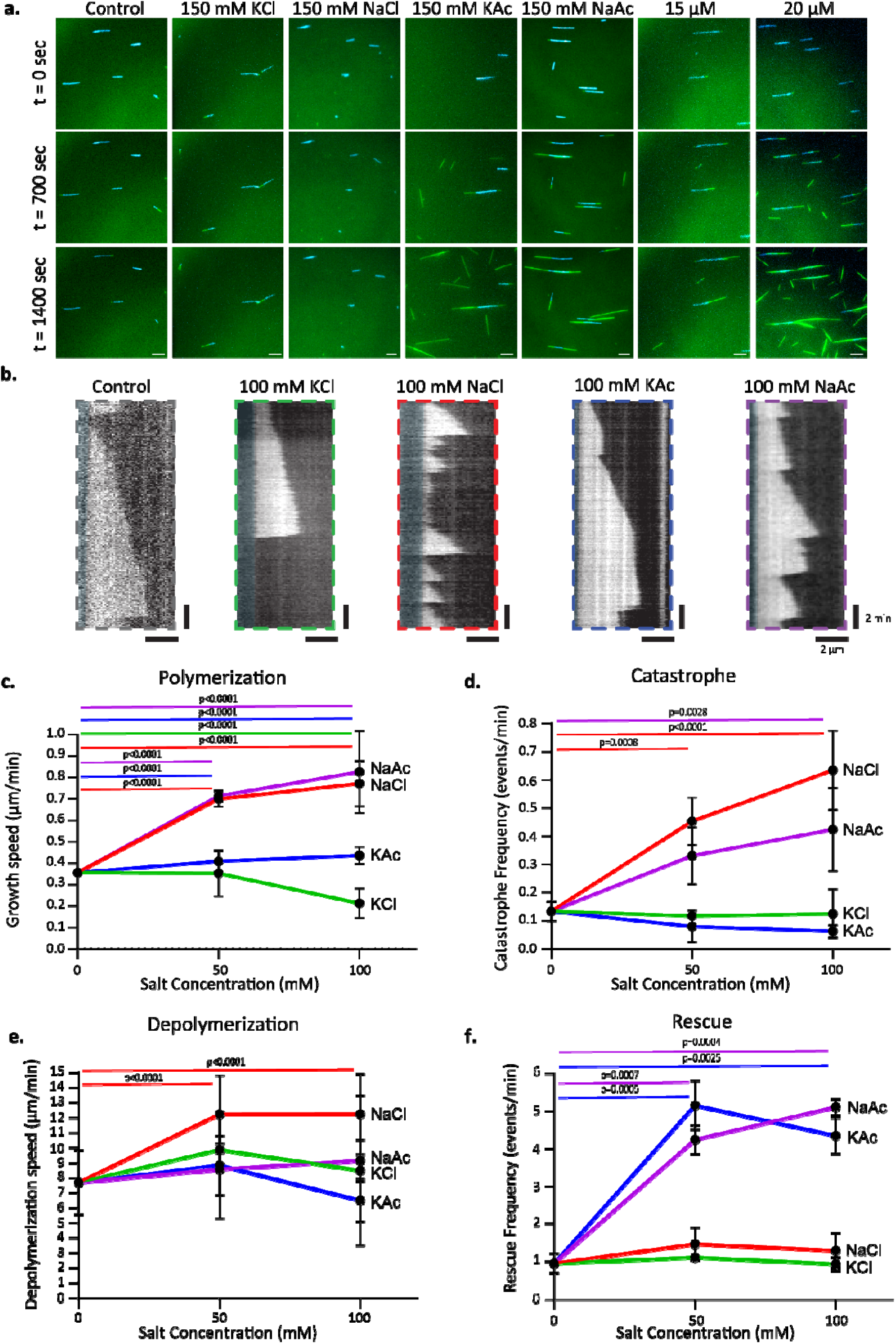
Ion effect on single microtubule dynamics. **a**. Representative images at three different time points (0, 700 and 1400 sec) of dynamic microtubule at 7 μM tubulin with 0 mM or 150 mM indicated salt. GMPCPP-stabilized microtubules (seeds) are in cyan, dynamic microtubules are in green. Scale bar: 5 μm. **b**. Representative kymographs of microtubules (7 μM tubulin). Control (0 mM salt, grey), 100 mM KCl (green), 100 mM NaCl (red), 100 mM KAc (blue) and 100 mM NaAc (purple) **c-f**. Mean microtubule growth speed (c), mean microtubule catastrophe frequency (d), mean microtubule depolymerization speed (e) and mean microtubule rescue frequency (f) (7 μM tubulin) with 0, 50, or 100 mM indicated salts. Experimental mean with SD, n = 3 independent experiments. Statistics: one-way ANOVA (the colored code was kept for statistics), the full statistic comparison between all conditions, individual and experimental mean values in **Sup. Fig. 1a-d**.

In our control condition (0 mM salt, in BRB80), 7 µM tubulin polymerized into microtubules at a speed of 0.36 ± 0.01 µm/min (**Fig. 2c**). Contrary to expectations from the bulk assay, microtubule growth speed increased only slightly with the addition of 50 mM or 100 mM KAc (**Fig. 2c**). Adding 100 mM KCl to the assay reduced the polymerization speed to 0.21 ± 0.07 µm/min (**Fig. 2c**). Conversely, the addition of either 50 mM NaCl or NaAc doubled the growth speed compared to the control (0.69 ± 0.03 and 0.71 ± 0.02 µm/min, respectively). Doubling the salt concentration to 100 mM only marginally further increased growth speed to 2.2-fold. However, at a concentration of 150 mM, both salts had contrasting effects: 150 mM NaCl completely inhibited polymerization, while 150 mM NaAc led to highly dynamic microtubules and to spontaneous nucleation (**Fig. 2a**). Collectively, these results show that salts containing the cation Na^+^ increase microtubule growth speed and indicate that the anion partners do not differentially affect the growth up to 100 mM but have a differential impact at higher concentrations (**Sup. Fig. 1a**).

We next studied how the different salts impact microtubule catastrophe events. Both KAc and KCl had no significant impact on catastrophe events (**Fig. 2d**). In contrast, the catastrophe frequency increased up to 5-fold in the presence of salts containing Na^+^, with NaCl amplifying the frequency nearly twice as much as NaAc **(Fig. 2d**). Combining 25 mM NaCl with 25 mM NaAc (50 mM Na^+^ in total) yielded comparable results to conditions with 50 mM NaCl alone (**Sup. Fig. 1b**), indicating that the increase in catastrophe events is caused by the Na^+^ cation. This is further supported by the results from a combination of 50 mM KAc and 50 mM NaAc (50 mM Na^+^ in total), showing a frequency comparable to that in the presence of 50 mM NaAc.

Upon catastrophe, the depolymerization speed was not altered in the presence of KAc, KCl nor NaAc, while 50 mM NaCl nearly doubled the depolymerization speed (12.24 ± 2.58 µm/min) compared to control 7.69 ± 2.15 µm/min (**Fig. 2e**). This result indicates that increasing shrinking velocities may depend on the cooperative effect between Na^+^ and Cl^-^. Doubling the NaCl concentration to 100 mM did not further increase the depolymerization speed. Microtubule depolymerization can occur in distinct phases with distinct speeds, though the net depolymerization speed remains constant (Luchniak et al., 2023). To test whether salts induce different depolymerization phases, we imaged at 1 second per frame. No alteration in depolymerization slopes was observed, indicating that depolymerization under the different conditions happened in a single regime (**Sup. Fig. 1f**).

Rescue events were rare, with a frequency of 0.95 ± 0.26 min^-1^ in control conditions and remained rare in the presence of KCl or NaCl (**Fig. 2f**). However, the rescue frequency increased up to 5-fold in the presence of Ac^-^ containing salts, reaching 5.11 ± 0.21^-1^ min for the highest concentration of 100 mM NaAc. The rescue-induced effect was observed even with a combination of 25 mM NaCl and 25 mM NaAc (total 25 mM Ac^-^), which tripled the rescue frequency compared to the control (**Sup. Fig. 1d**). Collectively, salts with the anion Ac^-^ boosted the rescue frequency, while those with the Cl^-^ anion did not significant affect rescue (**Fig. 2f**).

Taken together, 50 mM KCl did not have discernible impact on any of the microtubule dynamic parameters, explaining its preference in *in vitro* microtubule assays for modulating buffer ionic strength. The two main ions controlling dynamic are Na ^+^ and Ac^-^. While the cation Na^+^ boosts polymerization speed, it simultaneously is the major destabilizing factor by increasing the catastrophe frequency and the depolymerization speed, thereby increasing the dynamicity of microtubules. The presence of Cl^-^ might further potentiate Na^+^ effects. Conversely, the presence of Ac^-^ increases the rescue frequency.

### Acetate increases microtubule mass

Microtubules that undergo rescue are longer (Gardner et al., 2013; Schaer et al., 2023). Consistently, we measured a 2-fold increase in the mean microtubule length in the presence of 100 mM KAc compared to the control (**Fig. 3a, Fig. 2b and f**). The mean length of microtubules grown in the presence of NaAc is shorter but still 1.5-fold longer than the control (**Fig. 3a**). This likely results from a counter effect of Na^+^ and Ac-, while Ac^-^ induced rescues, Na^+^ caused catastrophe events (**Fig. 2d and f**). Consistent with the results that KCl only slightly reduced microtubule polymerization, the microtubule mean length was reduced slightly, while microtubules grown in the presence 100 mM NaCl where only one-third of the control length (**Fig. 3a**).

**Figure 3:**
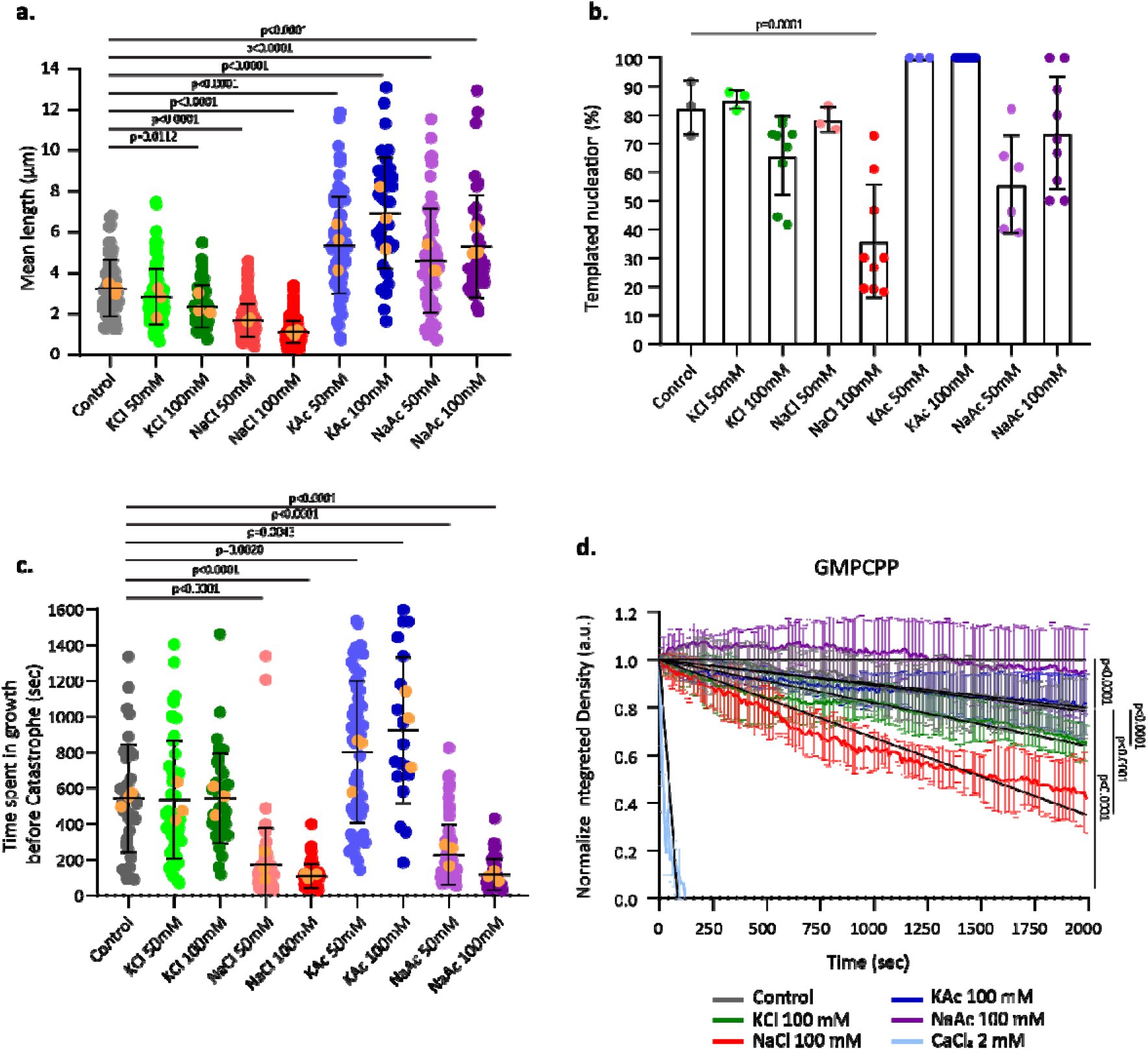
Ion effect on microtubule stability and nucleation. **a**. Mean microtubule length (7 μM tubulin) over 45 minutes with the indicated salt. The length was calculating from the cumulative length of single microtubules. Mean with SD, n = 3 independent experiments, individual values and experimental means (orange dots). **b**. Mean ratio of templated microtubule nucleation within 45 minutes (number of seeds with microtubule/ total number of seeds in the field of view). 7 μM tubulin, with the indicated salt. Experimental mean with SD, n = 3 to 10 independent experiments. **c**. Mean microtubule lifetime until the first catastrophe event, (7 μM tubulin) with the indicated salt. Mean with SD, n = 3 independent experiments, individual values and experimental means (orange dots). **d**. Normalized mean integrated fluorescent density of GMPCPP-microtubules (survival assay, see methods) was measured every 10 seconds over 200 frames. Experimental data are in colored lines and the linear regressions are in black (see methods). Mean with SD, n = 3 independent experiments. (a-d) Statistics: one-way ANOVA, for clarity only significant values are shown in the graph.

### Microtubule nucleation boosted by KAc

Templated nucleation of microtubule, nucleation from seeds, is kinetically unfavorable (Wieczorek et al., 2015). This is likely due to structural differences between the tapered dynamic microtubule ends and the double stabilized GMPCPP- (a slow hydrolysable analogue of GTP) Taxol-seeds, which tend to be blunter. Given that Ac-boosts spontaneous nucleation (**Fig. 1a and b**.), we asked whether Ac^-^ might promote templated microtubule nucleation. To address this, we measured the efficiency of seeds to nucleate microtubules in the presence of the different ions within 25 min (see Methods).

In average 80 % of the seeds polymerized microtubules in the control conditions, a ratio unchanged by the addition of either 50 mM KCl or 50 mM NaCl to the assay (**Fig. 3b**). However, increasing the concentration to 100 mM salt reduced microtubule formation by 1.25-fold (KCl) and 2.3-fold (NaCl). Thus, NaCl has the dual effect of enhancing polymerization speed while hindering new microtubule growth from seeds. This is further supported by the absence of microtubule polymerization beyond 100 mM NaCl (**Fig. 1a**).

In contrast to NaCl, the addition of 50 mM KAc to the reaction boosted templated microtubule nucleation to 100 %. With 100 mM Kac, we observed not only microtubule polymerization from each single seed, but also spontaneous nucleation in the bulk – a phenomenon that we observed only above 15 µM tubulin in the absence of KAc (**Fig. 1a**). At 150 mM KAc and 7 µM tubulin spontaneous nucleation was so pronounced that analysis of dynamic parameters became unfeasible due to the dense network, a condition resembling what we achieved at 20 µM tubulin in absence of KAc (**Fig. 1a**). The efficiency of the microtubule formation in the presence of 50 mM NaAc was reduced compared to the control condition but increased with increasing NaAc concentration (**Fig. 3b and Fig. 1a**). This result underscores a potential counteractive interplay between the anion Ac^-^ and the cation Na^+^, where increasing concentrations of Ac^-^ drive both templated and de novo nucleation even in presence of Na^+^.

### Microtubule aging accelerated by Sodium

The probability of a microtubule to undergo catastrophe increases with its lifetime (Duellberg et al., 2016b; Gardner et al., 2011; Odde and Buettner, 1995). This is due to the growing GTP-cap becoming more irregular and tapered over time, leading to a reduction in the GTP-density, which triggers catastrophe events (Duellberg et al., 2016b, 2016a). To explore how ions influence this aging process, we studied their impact on the growth-span of microtubules before the first catastrophe event occurred. The presence of KCl did not impact microtubule growth-span, whereas the presence of 100 mM NaCl reduced the growth-span by 6-fold compared to the control (**Fig. 3c**). Despite the general destabilizing effect of ions on microtubules (Bonacker et al., 2005; Perchellet et al., 1999; Shevtsov et al., 2016), KAc nearly doubled the growth-span relative to the control. Based on previously published EM data (Coombes et al., 2013) and theoretical models (Duellberg et al., 2016b), our results would imply that NaCl extends the tapper length of the GTP-cap, while KAc leads to a blunter end.

### Impact of nucleotide on salt driven disassembly

To address the impact of salts on microtubule stability, we monitored depolymerization of GMPCPP-stabilized microtubules in the absence of free tubulin (**Fig. 3d**). These GMPCPP-microtubules, which have more uniformly blunt ends compared to the variably aged ends of dynamic GTP-microtubules, provided: i) a more consistent environment to study how salts affect dimer removal, ii) a slower depolymerization that allows to capture fast processes that might go unnoticed during rapid depolymerization of GDP-microtubules, and iii) a different nucleotide (**Fig. 3d)**. The salt CaCl_2_ rapidly depolymerizes GMPCPP-microtubules (Müller-Reichert et al., 1998). We confirm this and observed that 2 mM CaCl_2_ depolymerized all GMPCPP-microtubules within 100 sec (**Fig. 3d**). The effects of the major salts used in this study were less pronounced; notably, 100 mM NaCl was the most effective destabilizer, reducing the microtubule mass by 50 % within 2000 seconds, consistent with the effect of NaCl on the mean microtubule length (Fig. 3a). The presence of 100 mM KCl slightly enhanced GMPCPP-microtubule disassembly. Surprisingly, 100 mM NaAc maintained the microtubule polymer compared to the control, indicating a stabilizing effect by either acting on the tip structure or on GMPCPP hydrolysis.

### Ions effect on tubulin-tubulin interaction in a GDP-lattice

The various salts distinctly influenced microtubule dynamics. To distinguish whether these effects came from alterations in the GTP-hydrolysis rate or from changes in tubulin-tubulin interactions, we study the impact of salts on a homogenous GDP-lattice, specifically the microtubule shaft (**Fig. 4a**). We end-stabilized GDP-microtubules and performed a damage assay in absence of free tubulin to monitor the microtubule breaking process over 35 minutes (**Fig. 4b**). KCl, which did not affect microtubule tip-dynamics, also showed no significant differences in shaft-dynamics compared to the control (**Fig. 4c**). However, with NaCl, 100 % of the microtubule broke, compared to 85 % in the control condition (**Fig. 4c**). Conversely, only 30 % of microtubules broke when incubated with KAc. Microtubule breaking is reduced to 70 % when incubated with NaAc, reinforcing our observation of the contrasting effect between Ac^-^ and Na^+^. This result shows that NaCl, NaAc and KAc act on tubulin-tubulin interactions and not on the hydrolysis rate. Like the case at the tip, at the microtubule shaft Na^+^ does destabilize the interactions and Ac^-^ has a stabilizing effect.

**Figure 4:**
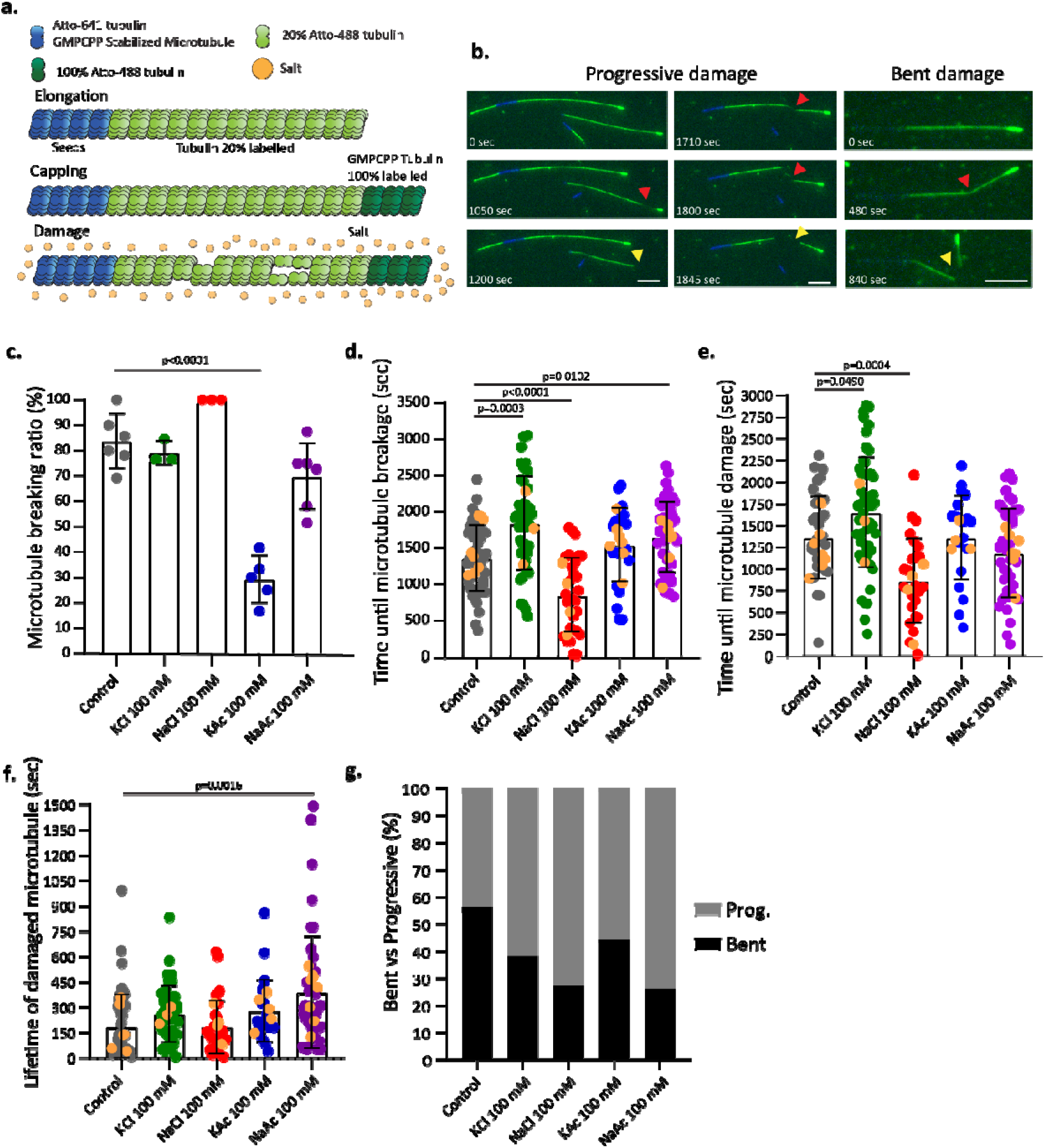
Ion effect on microtubule damage. **a**. Scheme of the damage assay. Salts were added during the damage step. **b**. Representative TIRF-images of microtubules showing progressive and bent damage over time. The first two columns show the same microtubules. Red arrowhead: damage site; yellow arrowhead: breakage of microtubule followed by depolymerization. Scale bar: 5 μm. **c**. Mean microtubule breaking ratio (number of broken microtubules/the total number of microtubules). **d**. Mean time until microtubule breakage (with the indicated salt), from the beginning of the assay until microtubule started to depolymerize, as indicated by the yellow arrowhead in Fig. 4b. **e**. Mean time until microtubule damage (with the indicated salt), from the beginning of the assay until the first sign of damage. **f**. Mean lifetime of damaged microtubules (seconds until microtubule depolymerized – seconds of first damage sign), with the indicated salt. (**d-f**) Experimental mean with SD, n = at least 3 independent experiments, individual values and experimental mean values (orange dots). Statistics: one-way ANOVA. **g**. Ratio between the two different damaged types with indicated 100 mM salt or 0 mM (Ctrl).

### The damage process of microtubule is affected by ions

To delve deeper into the breaking dynamics of microtubules in the presence of different salts, we measured the time it took for a microtubule to break (**Fig. 4d**). Since microtubules rapidly depolymerize post-breakage (Duellberg et al., 2016b; Tang-Schomer et al., 2010), we defined the breakage event as the moment when the microtubule began to depolymerize (yellow arrowhead **Fig. 4b)**. These results showed that microtubule break much faster in the presence of NaCl compared to the control, reducing the microtubule’s lifetime by 37 % (**Fig. 4d**). Surprisingly, while microtubules in the presence of KAc and NaAc had lifetimes comparable to the control, those incubated with KCl had a 1.3-fold increase in lifetime before breakage occurred, indicating a protective effect of KCl against breakage. Consistent with this, we observe a delay in the damage formation in presence of KCl (**Fig. 4e)**.

Although NaCl induced faster damage formation compared to the control (**Fig. 4e**), the duration from when the damage event occurred to when the microtubule ultimately broke was the same in both conditions (**Fig. 4f**). This was also the case for KCl and KAc, although the values are slightly higher. Only microtubules incubated with 100 mM NaAc were significantly more stable in this transition state, damage initiation to breakage, increasing this transition time by up to 2.1-fold (**Fig. 4f**). These results imply that although NaCl can initiate dimer dissociation from the shaft (**Fig 4c-g**), it does not influence the subsequent lateral extension of damage leading to breakage. This points to a unique impact of salts on the longitudinal versus lateral tubulin interactions within the lattice.

### Impact of salts on lateral and longitudinal tubulin-tubulin interactions

To study the potential impact of salt on these tubulin-tubulin interactions, we categorized the observed damages into two types: i) progressive damage, marked by a longitudinal extending decrease in fluorescent intensity that result in mechanical instable stretches that were wavering in and out of the field of view (see Movie, **Fig. 4b)** and ii) bent damage, characterized by an over 170° angle bending of the microtubule before the breakage, likely due to more lateral spreading of the damage **(Fig. 4b)**. The time required of these two damage types to initiate was comparable (red arrowhead in **Fig. 4a, Sup. Fig. 2a**). In the control condition both damage types occurred with equal probability (**Fig 4g**). However, the addition of salts shifted the likelihood towards progressive damage, with a maximum for NaCl and NaAc that had ¾ of the damage in the progressive type **(Fig. 4g**). These results support the idea where salts weaken lateral interactions allowing protofilaments to peel out of the lattice.

### Incorporation of GTP-tubulin into the microtubule shaft is altered by the ionic composition

We next studied whether the dynamics at the growing tip are differentially affected by salts compared to the more defined and rigid environment at the microtubule shaft. To do so, we performed our breakage assay in the presence of free GTP-tubulin, enabling the repair of damage sites (incorporation assay) and studied the impact of 100 mM salt on microtubule shaft dynamics (**Fig. 5a and b**). Consistent with our data from the breakage assay, in the presence of NaCl the number of incorporation events per µm increased by 2.5-fold compared to the control (**Fig. 5c**). Similarly, in the presence of NaAc the incorporation density increased by 1.8-fold. No significant changes were observed with either KAc or KCl. The presence of Na^+^ salts uniquely led to an increase in the number of repaired damages.

**Figure 5:**
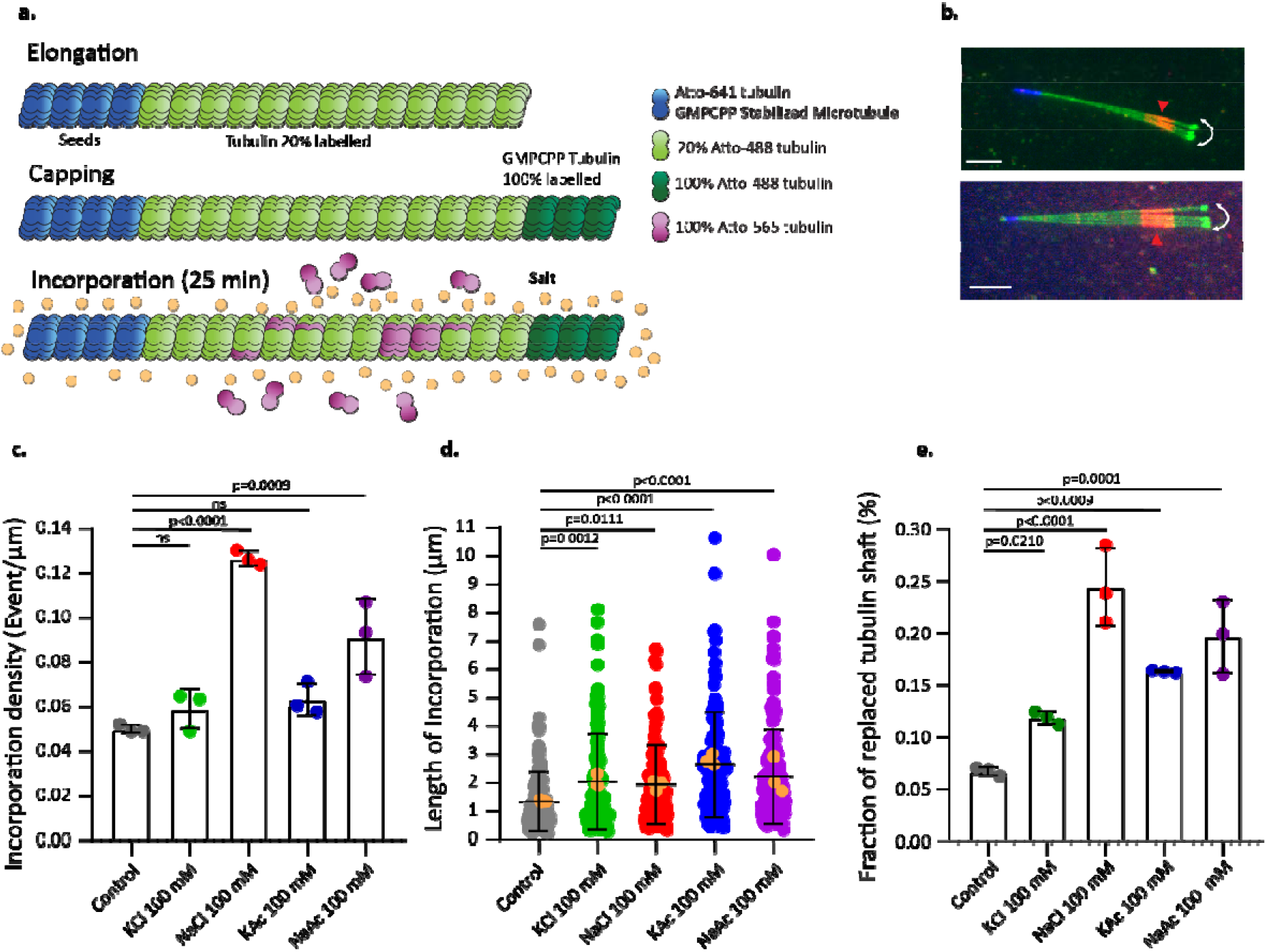
Ion effect on microtubule shaft dynamic. **a**. Scheme of the incorporation assay. Salts were added during the incorporation step for 25 minutes. **b**. Representative time projection (max intensity) of repaired microtubules after the incorporation assay. Scale bar: 5 μm. **c**. Mean incorporation density (event/μm) with the indicated 100 mM salt or 0 mM (control). Experimental mean with SD, n = 3 independent experiments. **d**. Mean length of incorporation sites with indicated salt. Experimental mean with SD, n = 3 independent experiments, individual values and experimental mean values (orange dots). **e**. Mean fraction of replaced tubulin shaft (total length of incorporation sites/total microtubule length) with indicated salt. Experimental mean with SD, n = 3 independent experiments. (**c-e**) Statistics: one-way ANOVA.

Furthermore, all salts tested increased the mean length of the incorporation sites, reaching a 2-fold increase with KAc compared to the control (**Fig. 5d**). This supports our observation that salts prompt progressive damage. These results indicate distinct effects of salts on both the removal and incorporation of tubulin dimers at damage sites. While KAc had no effect on the number of damage events, it led to the longest incorporation stretches, suggesting either increased longitudinal removal of dimers once damage occurs or higher repair efficiency.

The total fraction of exchanged tubulin along the shaft was highest in presence of NaCl, followed by NaAc, at 4-fold and 3-fold respectively, compared to the control (**Fig. 5e**). Thus, the number of events was more influential than the net repair efficiency, with KAc showing a 2.6-fold increase. Taken together, the microtubule incorporation assay revealed that the dynamics at both the growing microtubule tip and the shaft are regulated by the ionic composition surrounding the microtubule.

## DISCUSSION

Altogether, our results show that monovalent ions significantly affect microtubule dynamics. Using dynamic in vitro assays and TIRF microscopy, we precisely revealed how specific ions affect microtubule dynamics. We observed a strong effect of Na^+^ and Ac^-^ on microtubule stability. Interestingly, some dynamic parameters, such as rescue frequency, polymerization, and depolymerization speed, seemed to have already reached maximum changes at 50 mM salt, while catastrophe frequency continued to increase with higher salt concentrations. This indicates that different dynamic parameters may be governed by distinct electrostatic interactions or ionic interactions. Notably, 100 mM NaCl affects microtubule dynamics similarly to proteins that interact with microtubule ends, such as End Binding Proteins (EBs), thereby modulating microtubule dynamics (Bieling et al., 2008; Komarova et al., 2002; Miesch et al., 2023; Montenegro Gouveia et al., 2010; Vitre et al., 2008).

Throughout this study, Na^+^ demonstrated a pronounced destabilizing effect on the microtubule structure. Unlike a previous study using a combination of Taxol and NaCl (Wolff et al., 1996), our data show that net microtubule mass formation is not favored in the presence of Na^+^. This cation increases the catastrophe frequency, leading to shorter microtubules with reduced lifetimes (Fig. 2d,

Fig. 3a-3c). Similar to Ca^2+^, Na^+^ induces microtubule depolymerization (O’Brien et al., 1997; Weisenberg and Deery, 1981). However, the required concentration of Ca^2+^ to efficiently disassemble the microtubule structure is about 1000 times lower than that required for Na^+^ (0.1 mM and 100 mM, respectively) (Berkowitz and Wolff, 1981). Ca^2+^ is proposed to induce depolymerization by interacting with the C-terminal tail of tubulin (Lefèvre et al., 2011; Saoudi et al., 1995), while simulations predicted that Na^+^ is present both at the surface and inside the microtubule lumen (Shen and Guo, 2018). Whether Na^+^ binds to specific regions associated with tubulin-tubulin interaction remains elusive. Based on our damage and incorporation assay (**Fig**. 4 and **Fig**. 5), we hypothesize that Na^+^ weakens lateral interactions between tubulin heterodimers. This weakening likely leads to a higher catastrophe frequency (**Fig. 2c and 2d**), as MAPs that disrupt lateral interactions also cause microtubule depolymerization (Gardner et al., 2011), which is similarly the case when lateral interactions are weakened due to GTP hydrolysis (Akhmanova and Steinmetz, 2015). These weaker tubulin-interactions could furthermore explain the increase in microtubule damage, resulting in either more repair or breakage **(Fig. 4 and Fig. 5**).

In contrast to Na^+^, Ac^-^ promotes microtubule formation and stability. The rescue frequency is strongly increased in the presence of Ac^-^ (**Fig. 2f**), leading to longer microtubules with increased lifetimes (**Fig 3a and 3c**). However, this increase in rescue frequency was not caused by an increase in incorporation sites along the microtubule lattice (**Fig. 5c**). The higher incorporation efficiency of KAc and NaAc might be linked to their ability to trigger the highest rescue frequencies. However, NaCl, which had the highest number of incorporation sites, did not influence the rescue frequency. Thus, the increase in rescue frequency observed with KAc and NaAc likely stems from effects at the dynamic end of the microtubule, rather than from changes in shaft dynamics. We hypothesize that Ac^-^ stabilizes longitudinal interactions between tubulin heterodimers once integrated into the microtubule structure. This enhancement of interactions is further supported by the increase in nucleation efficiency (**Fig. 1a and 1b**), as longitudinal interactions are crucial for the nucleation process (Caudron et al., 2002; Job et al., 2003). Furthermore, the damage assay revealed a protective role of Ac against microtubule breakage during the assay (**Fig. 4c**), while Ac^-^ led to the longest incorporation stretches (**Fig. 5d**). In summary, in the presence of Ac^-^, microtubules break less but exhibit the highest incorporation length compared to other conditions (**Fig. 5c-e**), indicating that Ac^-^ enhances repair efficiency.

Our data suggests that, Ac^-^ counteracts the effects of Na^+^, while the Cl^-^ anion might synergize with Na^+^, leading to increased polymerization and depolymerization speeds in the presence of NaCl. The observed increase in repair sites with NaAc, alongside reduced breakage, reflects the opposing actions of Na^+^ in inducing damage and Ac^-^ in stabilizing damaged microtubules, preventing breakage.

Furthermore, the interplay between the two ions increased microtubule length, increased the rescue frequency, and enhanced net microtubule formation. This is especially pronounced at higher concentrations where 150 mM KAc nucleated microtubules, NaAc mildly nucleated microtubules and no microtubules were observed for NaCl. However, in terms of microtubule polymerization speed and catastrophe frequency, the effects of Na^+^ predominated over those of Ac ^-^. Further studies into the molecular mechanisms by which salts, particularly Na^+^ and Ac^-^, affect distinct microtubule dynamic parameters at the protein interaction level would be insightful.

In conclusion, the identity of the ion plays a more predominant role in the modulation of microtubule dynamics than its associated charge. While Na^+^ destabilizes the microtubule structure and Ac^-^ boosts microtubule formation, K^+^ and Cl^-^ did not show any drastic effect on microtubule dynamics. Therefore, the ionic environment in which microtubules grow must be considered according to the specific research question being addressed.

## ACKNOWLEDGEMENTS

We thank M. Andreu-Carbó and Pei-Tzu Yang for carefully reading our manuscript. ChatGPT for spelling and punctation correction.

## CONTRIBUTIONS

SF and CA conceptualized the project. SF performed and analyzed the *in vitro* dynamic assays, sedimentation experiments, incorporation, and breakage assay under the supervision of CA. SF and CA wrote the manuscript.

## MATERIAL AND METHODS

### Tubulin purification from bovine brain and tubulin labelling

Tubulin was purified from fresh bovine brain by two cycles of polymerization and depolymerization as previously described (Castoldi and Popov, 2003). A first polymerization-depolymerization cycle was performed in High-Molarity PIPES buffer (1 M PIPES-KOH at pH 6.9, 10 mM MgCl_2_, 20 mM EGTA, 1.5 mM ATP and 0.5 mM GTP) supplemented with 1:1 glycerol and depolymerization buffer (50 mM MES-HCl at pH 6.6 and 1 mM CaCl_2_) respectively. A second polymerization-depolymerization cycle was then performed: polymerization in High-Molarity PIPES buffer and depolymerization in 0.25XBRB80 complete after 15 min with 5XBRB80 to reach 1XBRB80 (80 mM PIPES at pH 6.8, 1 mM MgCl2 and 1 mM EGTA) respectively.

Labelled tubulin with ATTO-488, ATTO-565, or ATTO-647 (ATTO-TEC GmbH) and biotinylated tubulin were prepared as previously described (Hyman et al., 1991) with slight modifications. Tubulin was polymerized in glycerol PB solution (80 mM PIPES-KOH at pH 6.8, 5 mM MgCl_2_, 1 mM EGTA, 1 mM GTP and 33 % (v/v) glycerol) for 30 min at 37°C and layered onto cushions of 0.1 M NaHEPES at pH 8.6, 1 mM MgCl_2_, 1 mM EGTA and 60 % (v/v) glycerol followed by centrifugation. The pellet was resuspended in resuspension buffer (0.1 M NaHEPES at pH 8.6, 1 mM MgCl_2_, 1 mM EGTA, 40 % (v/v) glycerol) and incubated 10 min at 37°C with 1/10 volume of 100 mM ATTO-488, -565, or -647 NHS-fluorochrome or incubated 20 min at 37°C with 2 mM biotin reagent. Labelled tubulin was sedimented onto cushions of BRB80 supplemented with 60 % glycerol and resuspended in BRB80. A second polymerization-depolymerization cycle was performed before use. The labeling ratio was 11 % for ATTO-488 and 13 % for ATTO-565.

### Microtubule seeds preparation

Microtubule seeds were prepared by mixing 20 % ATTO-647-labelled tubulin and 80 % biotinylated tubulin to have a final concentration of 10 μM in BRB80 with 0.5 mM GMPCPP. The solution was incubated at 37°C for 45 minutes. 1 μM Paclitaxel was added to the solution and incubated at 37°C for 30 minutes. The solution was then centrifuged at 50.000 rpm at 25°C for 15 minutes. The pellet was resuspended with BRB80 supplemented with 1 μM Paclitaxel and 0.5 mM GMPCPP. Seeds were aliquoted and stored in liquid nitrogen.

### Imaging

Microscopy images were taken with an Axio Observer Inverted TIRF microscope (Zeiss, 3i) and a Prime BSI (Photometrics). A 100X objective (Zeiss, Plan-Apochromat 100X/1.46 oil DIC (UV) VIS-IR) was used. SlideBook 6 X64 software was used to record time-lapse imaging. Microscope stage conditions were controlled with the Chamlide Live Cell Instrument incubator (37°C).

### Flow chamber

Slides and coverslips were cleaned by two successive incubations and sonication: sonicated for 40 min in 1 M NaOH, rinsed in bidistilled water, sonicated in ethanol (96 %) for 30 min and rinsed in bidistilled water. Stocks of tri-ethoxy-silane-PEG-biotin and tri-ethoxy-silane-PEG were prepared at 1 mg/ml in 96 % ethanol and 0.02 % HCl. Slides and coverslips were dried with an air gun, placed into a Plasma cleaner (Electronic Diener, Plasma surface technology) for plasma treatment, followed by 2 days incubation with tri-ethoxy-silane-PEG (Creative PEGWorks) or a mixture of tri-ethoxy-silane-PEG-biotin and tri-ethoxy-silane-PEG with the ratio of 1:5 with gentle agitation at room temperature. Slides and coverslips were then washed in ethanol (96 %) and bidistilled water, dried with air gun and stored at 4°C. Flow chamber was assembled by fixing with double tap a tri-ethoxy-silane-PEG treated slide with a mixture of tri-ethoxy-silane-PEG-biotin and tri-ethoxy-silane-PEG treated coverslip with the ratio of 1:5.

### Microtubule dynamic assay

*In vitro* chamber was prepared by injecting successively 50 μg/mL neutravidin, BRB80, microtubule seeds (20 % ATTO-647-labelled tubulin and 80 % biotinylated tubulin) and washed with BRB80 to remove unattached seeds. 7 μM tubulin (20 % ATTO-488-labelled tubulin) in BRB80 supplemented with an anti-bleaching buffer (10 mM DTT, 0.3 mg/mL glucose, 0.1 mg/mL glucose oxidase, 0.02 mg/mL catalase, 0.125 % methyl cellulose (1500cP, Sigma) and 1 mM GTP) with the combination of 0 mM (control) or 50 mM or 100 mM salt or 150 mM (KCl, NaCl, KAc, NaAc) was injected in the chamber. The chamber was sealed for microscopy analysis. Microtubules polymerization was recorded over 300 frames at the interval of 5 seconds. High speed measurement was performed at 1 sec intervals, (**Sup. Fig. 1f**). Kymographs were generated and analyzed using an ImageJ macro. Polymerization and depolymerization speed were measured with the slope during the microtubule growing or shrinking phase respectively. Catastrophe frequency was calculated with the number of catastrophes spent during growth (frequency=catastrophe event/time in growth), Rescue frequency was calculated with the number of rescue event/ time spent in shrinkage. Microtubule mean length was measured with the cumulative length of single microtubule (from the first growth event of a microtubule until its full depolymerization). If a microtubule undergoes several catastrophe and rescue events before the full depolymerization, the measured mean length is the average of the different length that reach the microtubule before each catastrophe event. If a microtubule grows without catastrophe, the measured length is the final length that reached the microtubule during the acquisition. The mean ratio of templated microtubule nucleation was measured by the number of seeds with nucleated microtubule over the acquisition/total amount of seeds in the field of view. The microtubule lifetime is the time spent during the microtubule growth phase until the first catastrophe event. Graphs were created with GraphPad Prism8. One-way ANOVA was performed for statistics.

### Incorporation assay

Incorporation assay was adapted from the protocol published in Andreu-Carbó et al., 2022a. *In vitro* chamber was prepared by injecting successively 50 μg/mL neutravidin, BRB80, microtubule seeds (20 % ATTO-647-labelled tubulin and 80 % biotinylated tubulin) and washed with BRB80 to remove unattached seeds. 10 μM tubulin (20 % ATTO-488-labelled tubulin) in BRB80 supplemented with an anti-bleaching buffer (10 mM DTT, 0.3 mg/mL glucose, 0.1 mg/mL glucose oxidase, 0.02 mg/mL catalase, 0.125 % methyl cellulose (1500cP, Sigma) and 1 mM GTP) was perfused in the chamber. The chamber was incubated for 15 minutes at 37°C for microtubule polymerization from the seeds. A pre-warmed capping solution of 6 μM tubulin (100 % ATTO-488-labelled tubulin) in BRB80 supplemented with 0.5 mM GMPCPP was perfused in the chamber to exchange the polymerization buffer and then incubated for 15 minutes at 37°C. The chamber was washed twice with 100 μL pre-warmed BRB80 to remove unattached microtubules. An incorporation buffer containing 10 μM tubulin (100 % ATTO-565-labelled tubulin) in BRB80 supplemented with 0 mM (control) or 100 mM salt (KCl, NaCl, KAc, NaAc) was injected in the chamber and incubated at 37°C for 25 minutes. The chamber was washed with pre-warmed 100 μL BRB80 and an imaging solution of 6 μM tubulin (unlabelled) with BRB80 supplemented with anti-bleaching buffer was injected in the chamber. The chamber was sealed for microscopy analysis. Each position was recorded over 10 frames. Only fluctuating microtubules (**Fig. 5b**) were considered in the analysis to ensure that measured incorporation signal was not from an aggregate stacked on the surface. The mean incorporation density was calculated with the number of incorporation sites/ the total microtubule length. The mean fraction of replaced tubulin in the shaft was calculated with the total length of incorporation/ the total microtubule length. Measurements were done with Fiji. Graphs and statistical analysis (one-way ANOVA) were performed with GraphPad Prism8.

### Damage assay

Damage assay follows the same steps as the incorporation assay (explained above) until the capping solution incubation. The chamber was washed twice with 100 μL pre-warmed BRB80 to remove unattached microtubules. An imaging solution containing BRB80 supplemented with anti-bleaching buffer and 0 mM (control) or 100 mM salt (KCl, NaCl, KAc, NaAc) was perfused in the chamber. The chamber was sealed for microscopy imaging. Each position was recorded over 200 frames with an interval of 10 seconds. Only fluctuating microtubules were considered in the analysis to prevent any hypothetical glass stabilizing interaction with microtubules. The mean microtubule breaking ratio was calculated with the total number of broken microtubule/the total number of microtubules. The time required for breakage was measured from the beginning of the acquisition until the microtubule breakage. The time required to damage a microtubule was measured from the beginning of the acquisition until the first appearance of the damage sign. The mean lifetime of damaged microtubules was measured by subtracting the time required to breaks a microtubule with the time required to damage a microtubule. All the measurements were performed with Fiji. The graphs and the statistical analysis (one-way ANOVA) were performed with GraphPad Prism8.

### Turbidity assay

200 μL of 15 μM tubulin (unlabelled) were prepared in a BRB80 buffer supplemented with 1 mM GTP, 20 % glycerol (v/v) and 0 mM (control) or 50 mM or 100 mM salt (KCl, NaCl, KAc, NaAc). Solutions were transferred into a 96-well plate for turbidity analysis at 37°C. Absorbance was measured at 350 nm every 30 seconds over 3500 seconds with a Spectra MAX instrument. A control well (only BRB80) was included in the experiment to subtract the background. Data were extracted and processed with Microsoft Excel. A nonlinear regression (logistic fit; equation: Y=YM*Y0/((YM-Y0)*exp(-k*x)+Y0, where Y0 is the starting population (here Y0=0), YM is the maximal value (plateau) and k is the rate constant of the curve)), was applied on the graph to calculate the lag time, the plateau and the slope of the curve, **Fig. 1b**, with GraphPad Prism 8. One-way ANOVA was used to calculate statistics.

### Microtubule polymerization assay

10 μM tubulin (unlabelled) was polymerized in a BRB80 buffer supplemented with 1 mM GTP, 20 % glycerol (v/v), 0 mM (control) or 200 mM salt (NaCl or KAc) and incubated 30 minutes at 37°C. The solution was centrifuged at 50.000 rpm at 37°C for 20 minutes. The supernatant was harvested, and the pellet was resuspended in BRB80 buffer (similar volume as the starting volume). Samples were loaded in a SDS-PAGE gel followed by a Coomassie staining. Band intensities were measured on ImageJ and then plotted and analyzed with GraphPad Prsim 8. Two-way ANOVA was used to calculate statistics.

### GMPCPP Microtubule survival assay

10 μM tubulin (20 % labelled ATTO-488) was polymerized in a BRB80 buffer supplemented with 0.5 mM GMPCPP for 30 minutes at 37°C. GMPCPP microtubules were then mixed with anti-bleaching buffer (without GTP) and with 0 (Ctrl) or 2 mM CaCl_2_ or 100 mM of salt (KCl, NaCl, KAc and NaAC). Please note that EGTA was present in the BRB80, in the observed result at 2 mM CaCl_2_ it needs to be considered, that some of the Ca^2+^ was chelated by EGTA. 200 frames were recorded with an interval of 10 seconds. An arbitrary region with fixed size was randomly selected to measure the intensity over time for all experiments. Background intensity (selected region where no GMPCPP-microtubule were present) was subtracted from the measured integrated fluorescent intensity. The resulting value was normalized to 1 and the changes in fluorescent intensity were plotted over time. Measurements were performed with ImageJ and the graph plotted and statistical analysis (One-way ANOVA) were performed with GraphPad Prism8. Linear regressions were applied for each condition (Y=slope*x, Y0=1 as a restriction).

## SUPPLEMENTARY FIGURES

**Supplementary Figure 1:**
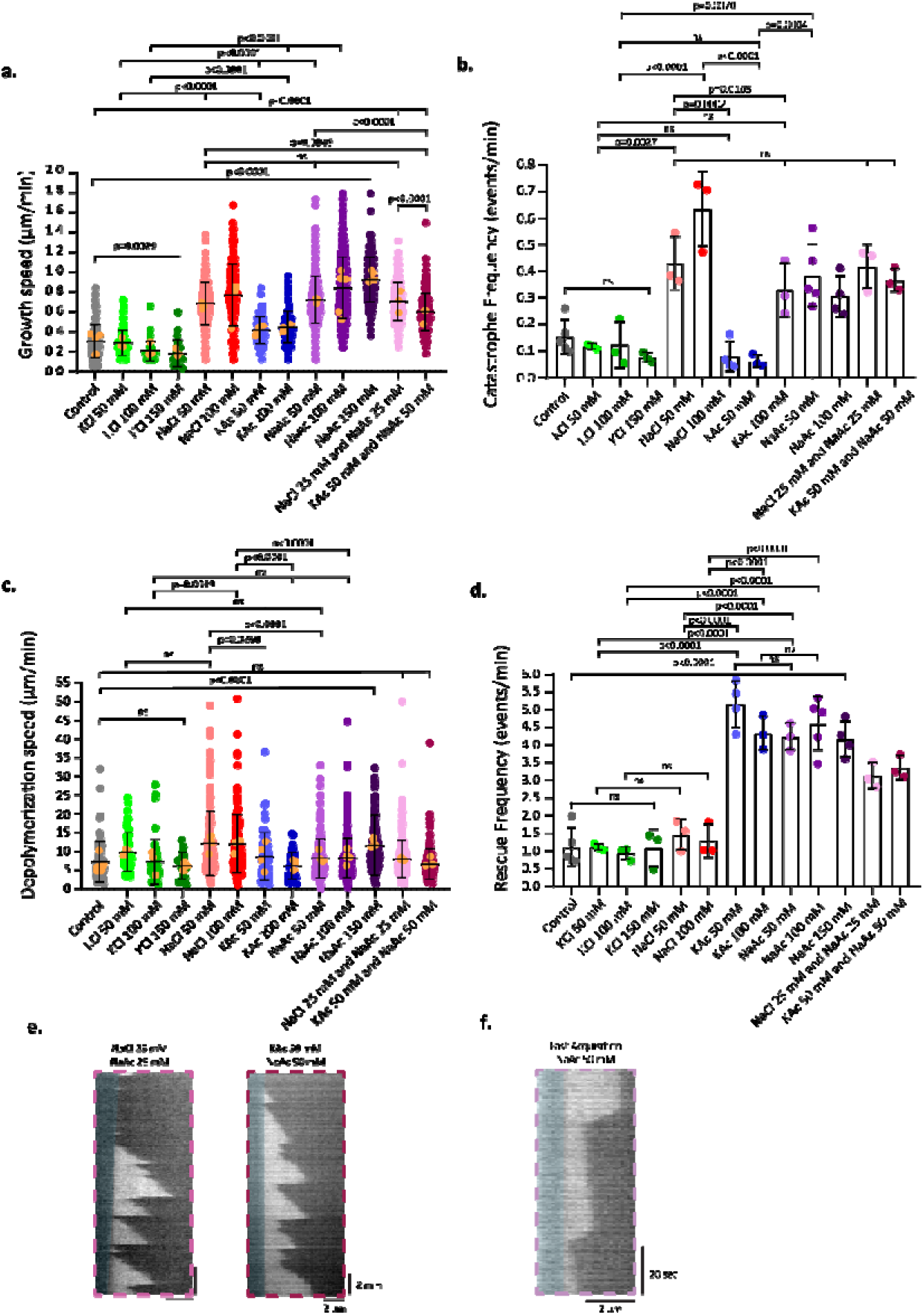
Single ion effect on microtubule dynamics. **a-d**. Mean microtubule growth speed (a), mean microtubule catastrophe frequency (b), mean microtubule depolymerization speed (c) and mean microtubule rescue frequency (d) (7 μM tubulin) with 0, 50, 100 or 150 mM indicated salts. Mean with SD, n = 3 to 5 independent experiments, individual values, and experimental means represented by orange dots for (a) and (c). Statistics: one-way ANOVA. **e and f**. Representative kymograph of microtubules (7 μM tubulin) with the indicated salt. The acquisition speed was 1 frame per second for (f).

**Supplementary Figure 2:**
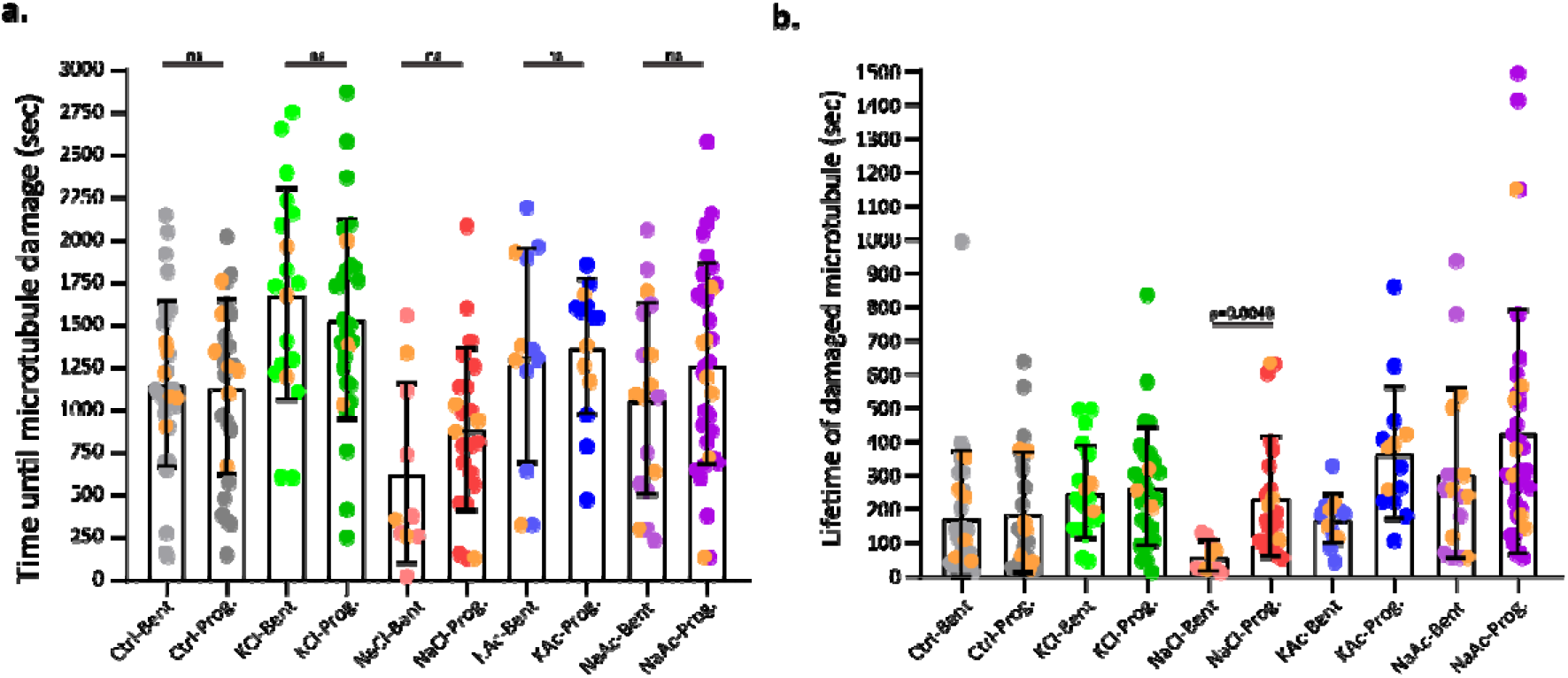
Ion effect on progressive and bent damages. **a**. Mean time until microtubule damage according to the damage type with the indicated salt. **b**. Mean damaged microtubule lifetime according to the damage type with the indicated salt. (a-b) Experimental mean with SD, n = at least 3 independent experiments, individual values and experimental mean values (orange dots). Statistics: one-way ANOVA.

